# Repeated Neuromuscular Electrical Stimulation Protocol Disrupts Muscle-to-Brain Signaling in Rats

**DOI:** 10.64898/2026.10.07.757107

**Authors:** Julien Wirtz, Yuan Wang, Rémi Chaney, Léa Wirtz, Quentin Drezet, Charalambos Papaxanthis, Aurore Quirié, Alexandre Méloux, Stéphanie Lemaire, Marina Cefis, Anne Prigent-Tessier, Philippe Garnier

## Abstract

**Objective:** Neuromuscular electrical stimulation (NMES) is increasingly used to preserve or restore muscle function in patients unable to exercise, and acute NMES improves cognitive performance and increases hippocampal mature brain-derived neurotrophic factor (BDNF). However, the consequences of repeated sessions for muscle-to-brain signaling remain poorly understood. We examined whether seven consecutive days of quadriceps NMES would reproduce this acute response.

**Methods:** Male Wistar rats received daily 30-min quadriceps NMES or SHAM sessions under isoflurane for seven consecutive days. Blood was collected immediately after the final session and 24 h later, when muscle and hippocampal tissues were also collected. An independent cohort underwent behavioral testing.

**Results:** Repeated NMES increased serum creatine kinase activity and quadriceps fibrillar collagen deposition without altering muscle mass, myofiber size, or degeneration-regeneration markers. These changes were accompanied by low-grade systemic inflammation and corticosterone elevation persisting 24 h after the final session. In the hippocampus, NMES increased inflammatory cytokine expression, astrocytic and microglial markers, and glucocorticoid receptor phosphorylation. Mature BDNF, TrkB phosphorylation and ERK1/2 signaling decreased, whereas proBDNF, p75NTR and JNK signaling increased; Akt and mTOR phosphorylation were unchanged. Synaptophysin, GAP43 and PSD95 also decreased. NMES increased marble burying and forced-swim immobility, without detectable changes in locomotion, recognition memory, or neuronal profile density.

**Conclusions:** Repeated daily NMES engaged a distinct, maladaptive mode of muscle-to-brain signaling rather than reproducing the benefits reported after acute stimulation. Stimulation parameters and recovery intervals that avoid this response need to be defined before repeated NMES can be recommended as an exercise substitute.

**Graphical abstract:** 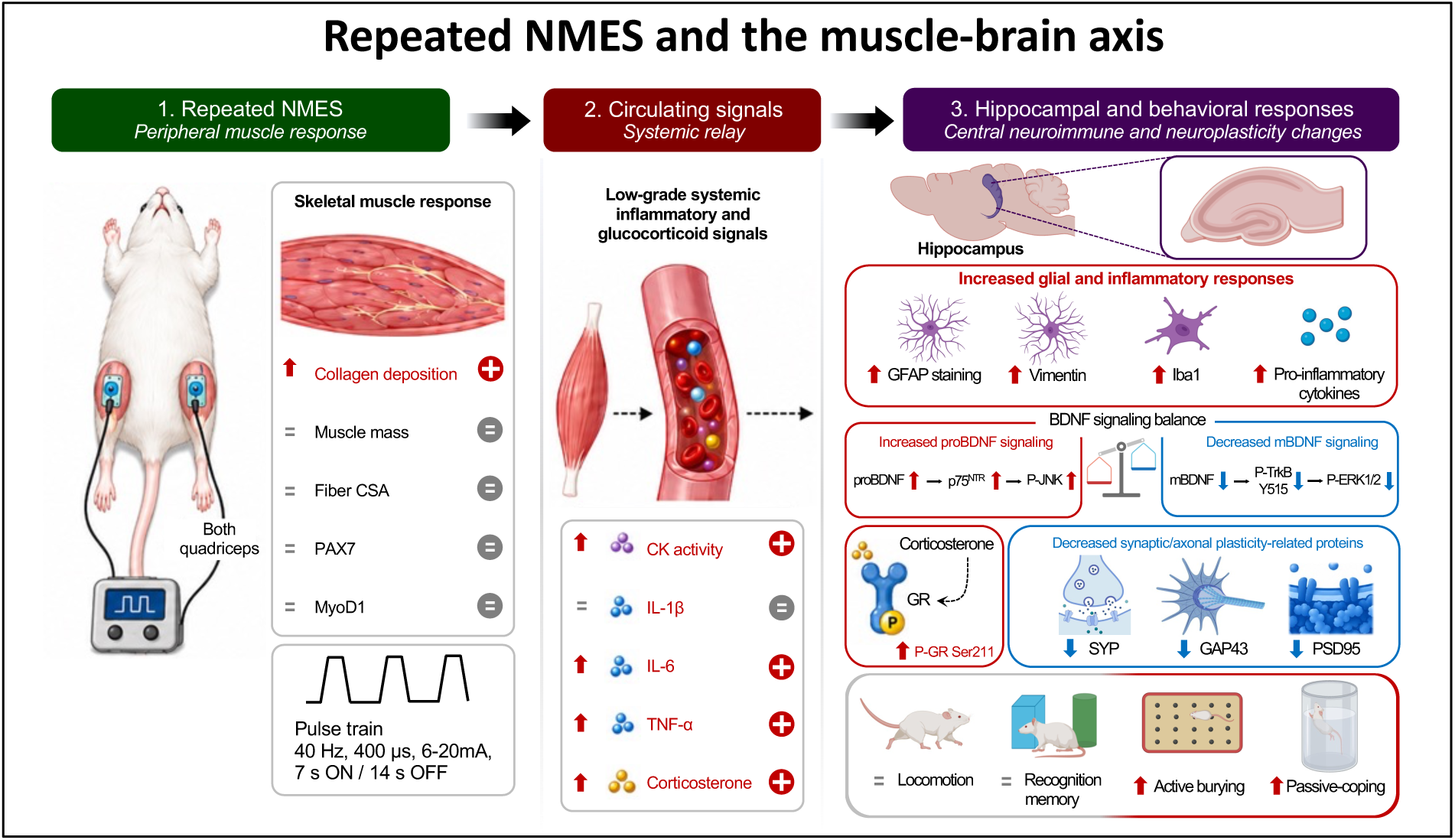

## 1. Introduction

Physical exercise is a multisystem challenge in which muscle contraction, cardiovascular recruitment, central motor command, and recovery interact to shape adaptations, including in the brain, where multiple mechanisms converge to increase BDNF (Cefis *et al*., 2023). Neuromuscular electrical stimulation (NMES) reproduces only part of this physiological sequence, since it generates forceful muscle contractions without requiring the voluntary motor command involved in conventional exercise. It therefore isolates the contracting muscle as a signal source, which makes it both clinically valuable and biologically informative. Clinically, it can preserve muscle function and limit disuse-related deconditioning in people whose capacity for voluntary contraction is reduced, whether by age, immobilization, or disease (Jones *et al*., 2016), and it induces genuine adaptations in the stimulated muscle, including hypertrophy and a fast-to-slow phenotype shift (Gondin *et al*., 2011), while potentially retaining some of the systemic benefits initiated by contracting skeletal muscle. Experimentally, electrically evoked contractions increase hippocampal BDNF in rodents (Maekawa *et al*., 2018) and improve cognitive performance in humans (Descollonges *et al*., 2024b, 2024a), supporting the possibility that NMES acts as a partial exercise mimetic rather than solely as a peripheral rehabilitation technique. We recently reported, in a combined human and rat study, that two 30-min quadriceps sessions delivered one week apart improved executive function and reduced anxiety in humans and increased hippocampal mature BDNF in rats, a protocol we defined as acute because each session was followed by a full week of recovery (Chaney *et al*., 2024).

However, biological equivalence between electrically induced contractions and voluntary exercise cannot be assumed. Electrically evoked contractions rely on a recruitment pattern that differs from that of voluntary exercise (Bickel *et al*., 2011; Enoka *et al*., 2020), and their systemic consequences therefore depend not only on the parameters of a given session but, critically, on how often sessions are repeated and on the recovery allowed between them. This becomes directly relevant when a stimulation paradigm comparable to one that is beneficial when applied with a week of recovery is instead delivered as daily 30-min sessions, a format chosen to mirror current physical activity recommendations of 150 to 300 min of moderate-intensity activity per week, commonly translated into a daily 30-min bout (Bull *et al*., 2020). Our initial expectation was that such repetition would not merely preserve the muscle-to-brain response observed after acute NMES but amplify it. Whether daily stimulation does so, or instead engages a qualitatively different response, remains unresolved. The question matters beyond NMES. Muscle-to-brain communication is generally presented as beneficial, with myokine signaling supporting hippocampal function (Pedersen, 2019). A muscle exposed to excessive or insufficiently recovered load, however, undergoes a maladaptive shift at the muscle level itself (Shorter *et al*., 2026), and may therefore signal to the brain through mediators that are inflammatory and neuroendocrine rather than trophic.

The hippocampus provides a sensitive readout of this duality, since its plasticity responds to exercise-related trophic signals and to stress mediators alike. Brain-derived neurotrophic factor (BDNF) is central to activity-dependent hippocampal plasticity, but its biological effect cannot be inferred from total BDNF expression alone. BDNF is first synthesized as the precursor proBDNF, which is then proteolytically cleaved into mature BDNF, and the two forms preferentially engage distinct receptors with opposing downstream consequences. Mature BDNF activates TrkB to support synaptic potentiation, whereas proBDNF signaling through p75^NTR^ facilitates long-term depression and can reduce dendritic complexity (Woo *et al*., 2005; Yang *et al*., 2014). The proBDNF to mature BDNF ratio is set downstream of transcription, by the rate of precursor synthesis and by the efficiency of its cleavage. This balance may determine whether the hippocampal environment remains permissive to structural and synaptic adaptation or instead favors synapse loss and dendritic retraction.

Two routes could account for such a shift. The first is neuroendocrine, since repeated peripheral challenge elevates circulating glucocorticoids and the hippocampus, which expresses glucocorticoid receptors at high density, is a principal target of the resulting corticosterone (McEwen & Akil, 2020). The second is inflammatory, because muscle activation that is intense or insufficiently recovered modifies the cytokine environment, and cytokines are potent modulators of hippocampal plasticity and of BDNF signaling in particular, acting in part through resident glia (Yirmiya & Goshen, 2011). Thus, when the contractile load imposed by NMES exceeds the adaptive capacity of the stimulated muscle, the resulting peripheral signal may differ in kind, and not only in magnitude, from that elicited by adaptive voluntary exercise. We previously showed that lumbar-nerve stimulation producing marked muscle damage alters hippocampal BDNF signaling and engages hippocampal glucocorticoid receptor signaling (Wirtz *et al*., 2025), establishing that damaging electrically evoked contraction can reach the hippocampus. Whether repeated stimulation does so in the absence of comparable muscle damage remains unknown.

The present study therefore examined whether daily quadriceps NMES, delivered as seven consecutive 30-min sessions, preserves an exercise-like muscle-to-brain response. We hypothesized that daily NMES would retain, and with repetition amplify, the hippocampal mature BDNF response previously observed after acute stimulation. Because repetition without recovery could instead engage stress-related signaling, muscle and circulating outcomes, the glucocorticoid response, affective and memory-related behavior, and hippocampal BDNF signaling and glial markers were assessed together.

## 2. Materials & methods

### 2.1. Animals

The experiments were conducted in accordance with the guidelines set forth by the French Department of Agriculture (license 21-CAE-102) and approved by the local ethics committee (Ethics Committee of Animal Experimentation, Dijon, registration number 105). Two independent cohorts of 8-week-old male Wistar rats, purchased from Janvier Labs (Le Genest Saint Isle, France), were used in this study. Animals were housed four per cage at 22 ± 2°C and 45–65% relative humidity, with ad libitum access to food and water and a 12-h light/dark cycle (lights off at 21:00, lights on at 09:00). Experimental procedures were performed between 06:00 and 09:00, during the last three hours of the dark phase, when the animals were at the end of their active period. After a 7-day quarantine period in the animal facility, rats underwent a further 7-day habituation period to handling by the experimenters. Animals were allocated to NMES or SHAM conditions by block randomization within each cage, with two NMES-treated and two SHAM animals housed together to minimize cage-related confounding. The first cohort was dedicated to molecular, biochemical, and histological analyses and included 24 animals assigned to SHAM (n = 12) or NMES (n = 12). Within this cohort, 8 animals per group were allocated to biochemical and molecular analyses and 3-4 per group to histological analyses, according to a predefined allocation. A second independent cohort was used for behavioral experiments and included 14 animals assigned to SHAM (n = 7) or NMES (n = 7). Group sizes were based on those used in previous studies applying the same stimulation model in our laboratory (Chaney *et al*., 2024; Wirtz *et al*., 2025); no a priori power analysis was performed. A priori exclusion criteria included visible injury, persistent abnormal behavior, overt signs of pain or distress, impaired locomotion, or any predefined humane endpoint requiring veterinary intervention. No animal met the predefined exclusion criteria. Tissue allocation between biochemical and histological analyses was predefined, and individual data points are shown for every animal in the figures. The experimental design is summarized in Fig. 1A.

**Figure 1.**
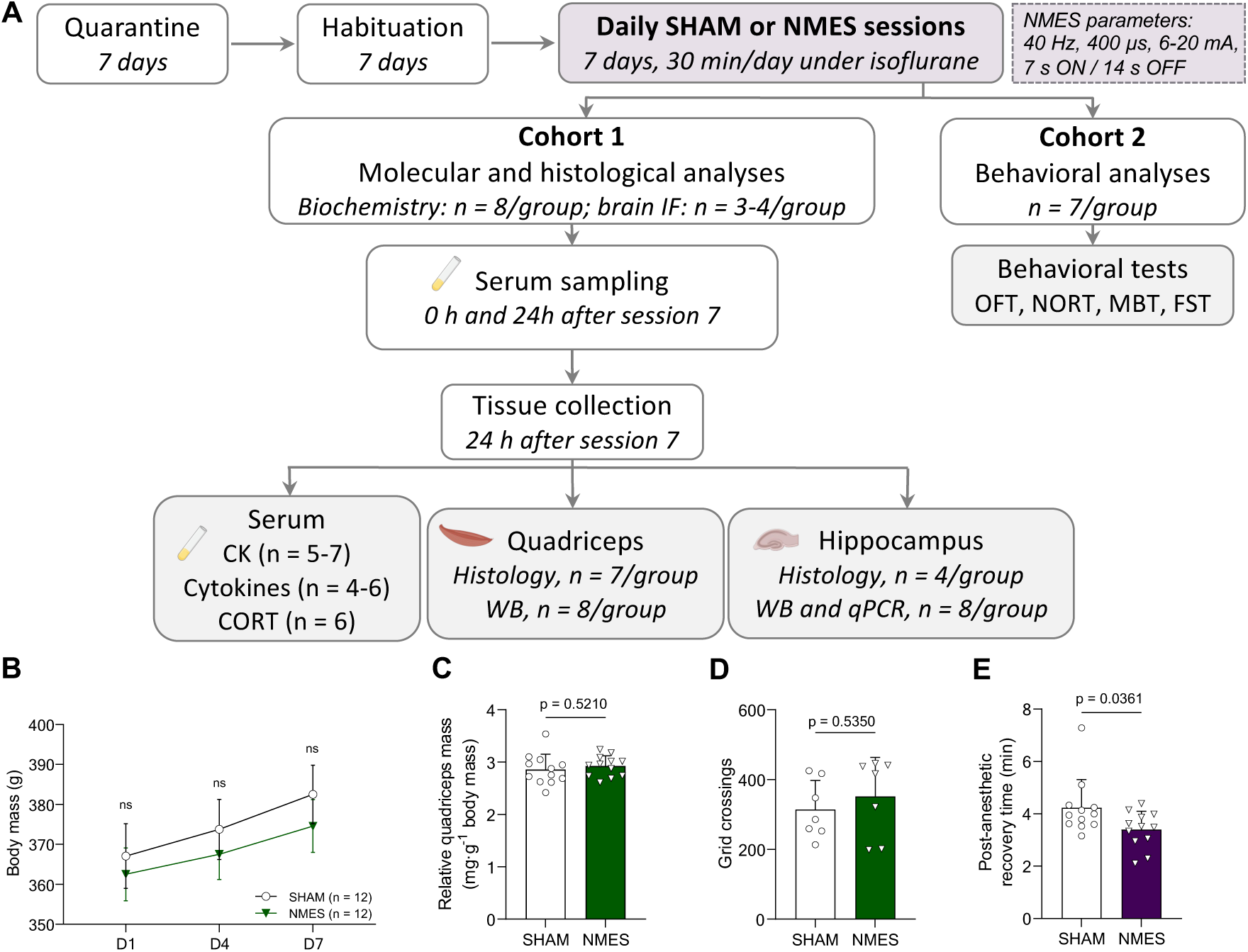
Experimental design and general physiological outcomes following repeated NMES. (A) Experimental design. Rats underwent 7 days of quarantine and 7 days of habituation, followed by 7 daily 30-min SHAM or NMES sessions under isoflurane. NMES was delivered at 40 Hz with 400 µs pulses, an intensity of 6-20 mA, and a 7-s ON/14-s OFF duty cycle. Cohort 1 was used for molecular and histological analyses, with blood collected immediately after session 7 (0 h) for creatine kinase activity, a second blood sample collected 24 h later for creatine kinase activity, corticosterone and cytokines and tissues collected at the same 24-h time point. Cohort 2 underwent behavioral testing. IF, immunofluorescence; OFT, open-field test; NORT, novel object recognition test; MBT, marble burying test; FST, forced swim test; CK, creatine kinase; CORT, corticosterone; WB, Western blot. (B) Body mass measured on days 1, 4, and 7 of the SHAM or NMES intervention period. (C) Quadriceps mass normalized to body mass. (D) Spontaneous locomotor activity assessed as the number of grid crossings, measured in the behavioral cohort. (E) Post-anesthetic recovery time following the final session. Data are presented as individual values with mean ± SD. ns, not significant. Longitudinal data were analyzed using two-way repeated-measures ANOVA with Šidák-adjusted comparisons, and single-time-point comparisons using the appropriate two-tailed unpaired test as described in the Methods.

### 2.2. NMES protocol

Following the 7-day handling habituation period, rats underwent daily 30-min NMES or SHAM sessions between 06:00 and 09:00 for seven consecutive days, as summarized in Fig. 1. Rats were anesthetized with isoflurane (4% for induction and 2% for maintenance through a standard rat nose cone). Anesthetic depth was assessed throughout the procedure by monitoring the absence of the pedal withdrawal reflex in response to toe pinch. Body temperature was maintained at approximately 37°C using a thermostatically controlled heating pad. Two self-adhesive square electrodes measuring 1 × 1 cm (Fyzéa, BAS95050, La Roche-Sur-Yon, France) were placed on the upper and lower parts of each quadriceps, respectively, to induce bilateral simultaneous muscle contraction. The electrodes were connected to a portable stimulator (Cefar Rehab X2, 111126, Lyon, France), and the stimulation parameters were as follows: biphasic current at 40 Hz, 400-µs pulse width, and cycles of 7 s ON and 14 s OFF for 30 min. Stimulation intensity was progressively increased from 6 to 20 mA during each session according to the same predefined sequence for all animals and sessions. Intensity was therefore not titrated to an individually determined evoked force, which was not recorded. This sequence produced a reproducible visible quadriceps contraction without overt whole-limb displacement or joint movement. Contractions were visually verified to be sustained and fused throughout each ON period. NMES-treated and SHAM animals underwent identical handling, anesthesia, electrode placement, session duration, and experimental timing, with electrical current delivered only in the NMES group. Following each session, animals were observed continuously during the initial recovery period and reassessed approximately 30 min later. Post-anesthetic recovery time was defined as the interval between discontinuation of isoflurane and the resumption of coordinated ambulation, which occurred after eye opening and recovery of the righting reflex. Monitoring included grooming, posture, locomotion, abnormal facial expressions, guarding or limb unloading, withdrawal or freezing behavior, and inspection of the stimulated skin for erythema or injury. No overt clinical signs of persistent discomfort, distress, abnormal gait, impaired movement, or skin damage were observed. Body mass was recorded throughout the protocol. Post-anesthetic recovery time was recorded after each session and averaged within each animal.

### 2.3 Experimental design and timeline

Two independent cohorts were used. The molecular cohort was dedicated to biochemical and histological analyses, whereas the behavioral cohort comprised seven animals per group and underwent a standardized sequence of behavioral tests ordered from the least to the most aversive paradigm. In the behavioral cohort, the habituation phase of the novel object recognition task (NORT), performed in the empty arena without objects, was conducted on Day 6, 2 h after the NMES or SHAM session. Locomotor activity was quantified during this session. On Day 7, 24 h after arena habituation and 2 h after the final NMES or SHAM session, animals underwent the NORT familiarization phase with two identical objects. The short-term memory test was performed 1 h later. On Day 8, 24 h after the final stimulation session, animals underwent the NORT long-term memory test, followed 2 h later by the marble-burying test and, after a further 2-h interval, by the forced-swim test. In the molecular cohort, blood was collected immediately after the seventh NMES or SHAM session (0 h) for serum creatine kinase assessment. A second blood sample was collected 24 h later for creatine kinase, corticosterone, and circulating cytokine measurements. Muscle and brain tissues were collected at the same 24-h time point.

### 2.4. Behavioral testing procedures

Behavioral experiments were conducted in a separate cohort. NMES-treated and SHAM animals were tested in paired succession under identical environmental conditions, within the same time window, and by the same experimenter. Testing order was counterbalanced between groups to minimize systematic effects of time or handling. Tests were administered in a fixed sequence progressing from the least disruptive to the most aversive procedure: open field, novel object recognition, marble burying, and forced swimming. The forced swim test was always performed last to prevent its aversive nature from influencing the other behavioral outcomes. Behavioral testing began at the end of the dark phase and continued into the early light phase, following an identical schedule for both groups. Animals were visually inspected before each test for abnormal posture, impaired gait, guarding, reduced spontaneous movement, abnormal facial expression, or other overt signs of discomfort. No animal required test postponement or removal on welfare grounds. Animals were transferred to the testing room at least 20 min before testing. All tests were performed in a quiet, temperature-controlled room under standardized lighting conditions of 20 lux. Apparatus and objects were cleaned with 70% ethanol between animals and allowed to dry completely before reuse. Behavioral scoring was performed offline by an experimenter blinded to group allocation.

#### 2.4.1 Open Field Test

Locomotor activity was assessed using the open field test. The apparatus consisted of a 60 × 60 cm square arena enclosed by 60-cm-high opaque walls. The floor was divided into 10 × 10 cm squares to facilitate manual scoring of locomotor behavior. Each animal was individually placed in a corner of the arena and allowed to explore freely for 10 min. The number of grid crossings (defined as the animal moving all four paws into a new square) was recorded as an index of spontaneous locomotor activity. Sessions were video-recorded and scored offline by a blinded experimenter. This session also served as the habituation phase of the novel object recognition test described in Section 2.4.2.

#### 2.4.2 Novel Object Recognition Test (NORT)

The NORT was used to evaluate recognition memory. The protocol consisted of three phases: habituation, familiarization and testing. On Day 6, animals were habituated to the arena for 10 min without any objects, in the session described in Section 2.4.1. On Day 7, during the familiarization phase, two identical objects (objects A/A’) were placed symmetrically in opposite corners of the arena, and rats were allowed to explore for 10 min. After a 1-h retention interval, the test phase assessing short-term memory was conducted by replacing one of the familiar objects with a novel one of similar size and texture (objects A/B). Rats were again allowed to explore for 10 min. On Day 8, long-term recognition memory was assessed by presenting one object previously encountered during familiarization together with a novel object. Rats were allowed to explore for 10 min. The identity and position of the retained familiar object and the novel object were counterbalanced across animals. Exploration time directed at each object was recorded, and the discrimination index was calculated as follows:

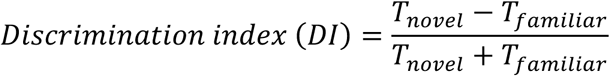

Object exploration was defined as the animal directing its nose toward the object at close distance and/or physically sniffing it. Climbing or sitting on the object was not scored as exploration. All sessions were video-recorded and scored offline by an experimenter blinded to group assignment. Total object exploration time was calculated as the sum of the time spent exploring both objects during each session. This measure was examined to verify comparable engagement with the task between groups. The objects used were previously validated according to a published protocol ensuring equivalent spontaneous exploratory value (Inayat *et al*., 2021).

#### 2.4.3. Marble burying test (MBT)

The MBT was used as an index of repetitive digging and defensive-like behavior. Each rat was individually placed in a standard cage (53 × 32 × 20 cm) filled with 5 cm of bedding material. Twenty dark blue glass marbles (15 mm diameter) were evenly spaced in a 4 × 5 grid on the surface. Rats were allowed to explore freely for 30 min. At the end of the session, the number of marbles buried at least two-thirds beneath the bedding was counted. Because marble burying may reflect overlapping defensive, anxiety-related, and repetitive dimensions, this outcome is reported as burying behavior rather than as a specific index of anxiety.

#### 2.4.4. Forced swim test (FST)

The FST was used to assess passive coping-related behavior. Each animal was placed individually in a transparent cylindrical tank (height: 1 m, diameter: 20 cm) filled with water rendered opaque with non-fat dry milk to a depth of 30 cm, preventing the animal from touching the bottom. Water temperature was verified before each test and maintained at 25 ± 1°C. The water was replaced between animals. Rats were gently placed into the tank for a 6-min test session. A single test session was used, without the pretest swim included in the classical two-day protocol, in order to avoid introducing an additional major stressor into a study designed to characterize the stress response itself. Animals were continuously monitored throughout the test. Immediately after the session, rats were gently dried with a clean towel and returned to their home cage, where they were kept under a heating lamp until fully dry and displaying normal spontaneous behavior. Forced-swim behavior was video-recorded and quantified offline by an experimenter blinded to group assignment. Immobility was defined as floating with only the movements necessary to keep the head above water and was used as an index of passive coping behavior.

### 2.5. Blood and tissue collection

Immediately after the final NMES or SHAM session, while animals remained under isoflurane anesthesia, blood was collected through a single incision of the warmed tail and transferred into serum-separation tubes. Hemostasis was achieved by applying gentle pressure to the incision site until bleeding had ceased. Blood samples were allowed to clot for 45 min at room temperature and then centrifuged at 2,000 × g for 15 min. Serum aliquots were stored at −80°C until analysis. These samples were used to measure circulating creatine kinase activity and therefore represented the immediate post-stimulation time point. Twenty-four hours after the final NMES or SHAM session, animals received buprenorphine analgesia (Buprecare, 0.05 mg/kg, intraperitoneally). Thirty minutes later, deep anesthesia was induced with ketamine/xylazine (75 mg/kg and 8 mg/kg, intraperitoneally). Adequate anesthetic depth was confirmed by the absence of the pedal withdrawal reflex. A second blood sample was collected by intracardiac puncture under terminal anesthesia, and processed for serum preparation to assess corticosterone, cytokines, and circulating CK activity at the 24-h recovery time point. The interval between anesthetic induction and blood collection was kept constant across animals and identical between groups. Terminal blood sampling was performed within the same daily time window as the stimulation sessions, so that samples from both groups were collected at a comparable circadian phase, near the diurnal nadir of corticosterone. Animals were then transcardially perfused with physiological saline before tissue collection. Quadriceps muscles were rapidly excised and divided, with one half snap-frozen in liquid nitrogen for biochemical analyses and the other half embedded in tragacanth gum (Sigma, G1128), frozen in precooled isopentane, and stored at −80°C for histological analyses. According to the predefined endpoint allocation, brains were either immersion-fixed in 4% paraformaldehyde for 48 h at room temperature for histological processing, or rapidly removed for hippocampal dissection. All brains were fixed and processed in parallel under identical conditions. Dissected hippocampi were snap-frozen in liquid nitrogen and stored at −80°C until biochemical and molecular analyses. Animals from the molecular and histological cohort were not exposed to behavioral testing.

### 2.6. Multiplex analyses

Serum IL-1β, IL-6, and TNF-α concentrations were measured in the samples collected 24 h after the final NMES or SHAM session, using MILLIPLEX magnetic bead panel kits (Merck Millipore, Burlingame, CA, USA). Samples were analyzed on a Luminex MAGPIX system (Luminex Corporation, Houston, TX, USA) using MILLIPLEX Analyst software (Millipore, St. Charles, MO, USA). All samples were assayed in duplicate, and a positive control was included to verify assay performance. The assay limit of detection was 1.22 pg·mL-1. Only IL-6 yielded values below this limit; these were classified as non-detectable and, for graphical representation only, were plotted at 1.22 pg·mL-1 without being interpreted as exact concentration measurements. The proportions of samples with detectable IL- 6 concentrations were compared between groups using a two-sided Fisher’s exact test.

### 2.7. Corticosterone measurement

Serum corticosterone was quantified by liquid chromatography–tandem mass spectrometry using a commercially available MassChrom® kit (Chromsystems, Germany) following the manufacturer’s instructions. A 200-µL aliquot of fivefold-diluted serum was mixed with an internal standard and loaded onto a solid-phase extraction (SPE) plate with an extraction buffer via centrifugation. After two washing steps and elution in 500 µL of elution buffer, the extracts were evaporated to dryness and reconstituted in 100 µL of reconstitution buffer. Chromatography was performed with an Acquity UPLC system (Waters Corporation, Milford, MA, USA) coupled to a Xevo TQ-S micro tandem mass spectrometer (Waters Corporation, Milford, MA, USA).

### 2.8. Creatine kinase activity measurement

Serum creatine kinase (CK) activity was measured immediately after the final session and again 24 h later using a CK-NAC IFCC assay (BIOLABO, Maizy, France; Ref. 92207), according to the manufacturer’s instructions. The assay is based on the standardized IFCC kinetic method, in which CK-catalyzed ATP production is coupled to hexokinase and glucose-6-phosphate dehydrogenase reactions, resulting in the formation of NADPH. The increase in absorbance at 340 nm is directly proportional to CK activity. CK activity was expressed as U·L-1 and calculated using the theoretical factor specified by the manufacturer for a 1-cm optical path length at 37°C: CK activity (U/L) = ΔAbs/min × 3333.

### 2.9. RNA extraction and RT-qPCR

Total RNA was extracted from hippocampal tissue using TRIzol reagent (Invitrogen) according to the manufacturer’s instructions. RNA concentration and purity were assessed using a NanoDrop™ spectrophotometer (Thermo Scientific). Complementary DNA (cDNA) synthesis was conducted with the iScript cDNA Synthesis Kit (Bio-Rad). Quantitative PCR was carried out with PowerUp SYBR Green Master Mix (Applied Biosystems, Life Technologies) on the StepOnePlus™ Real-Time PCR System (Applied Biosystems). Each sample was analyzed in technical duplicate, and relative gene expression was calculated using the 2−ΔΔCt method. Gene expression was normalized to the geometric mean of β-actin and 18S rRNA as reference genes. Reference-gene expression was stable across experimental groups. Primer sequences are detailed in Table 1.

**Table 1:**
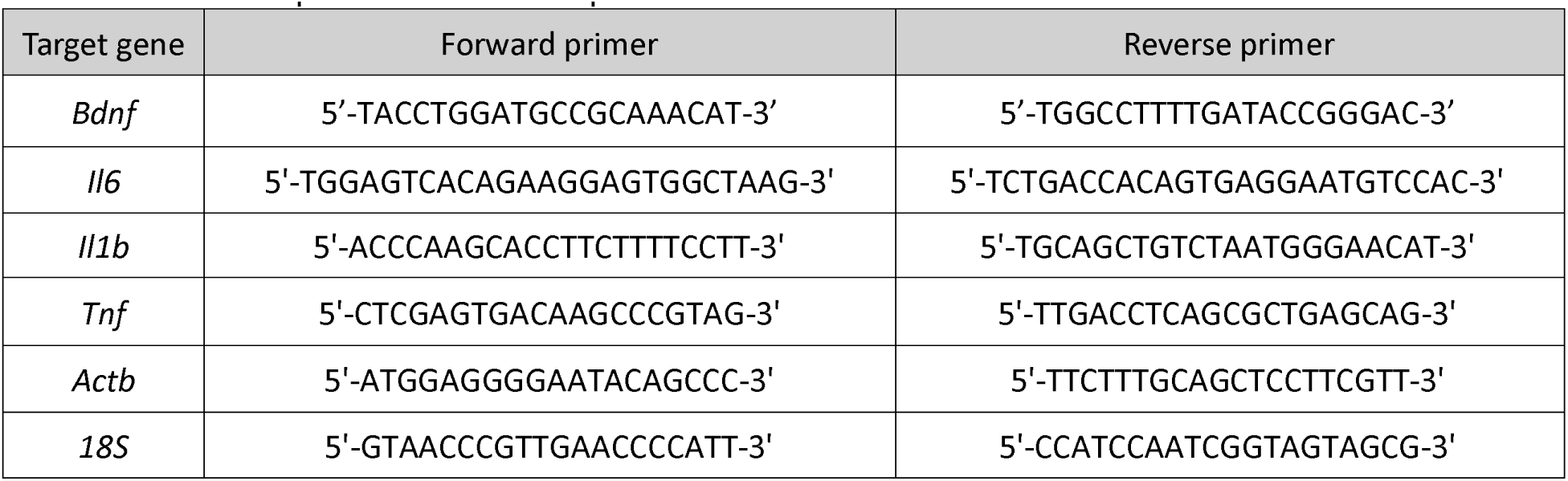
List of primers used for RT-qPCR.

### 2.10. Protein extraction and Western blotting

Frozen muscle and brain tissue samples were homogenized in a lysis buffer (100 mM Tris base, 150 mM NaCl, 1 mM EGTA, 1% Triton X-100, protease and phosphatase inhibitors, pH 7.4) using a 7-fold volume ratio in the Precellys system (Bertin Technologies). Homogenates were centrifuged at 12,000 × g for 10 min at 4°C. The supernatant containing total soluble proteins was collected, and protein concentration was determined using the Lowry assay (Pierce™ Modified Lowry Protein Assay Kit, Thermo Fisher Scientific). For each target, equal amounts of protein were loaded for all samples; the amount loaded ranged from 1 to 40 µg depending on target abundance and assay optimization. Samples were mixed with Laemmli buffer (Tris 125 mM, SDS 4%, glycerol 20%, bromophenol blue 0.01%, β-mercaptoethanol 5%), heated at 95°C for 10 min and separated by SDS-PAGE on 7.5% or 12% polyacrylamide gels (TGX Stain-Free FastCast Acrylamide Kit, 7.5%, 1610181; 12%, 161-0185, Bio-Rad). Gels containing proteins were activated under UV light with a ChemiDoc imaging system (12003153, Bio-Rad) using the “Stain-Free Gel Activation (45 s)” program. Proteins were then transferred onto 0.2 µm nitrocellulose membranes (1620112, Bio-Rad) using a semi-dry transfer system (TransBlot Turbo, 1704150, Bio-Rad). Membranes were blocked for 1 h at room temperature in 5% non-fat dry milk prepared in TBS-T (Tris-buffered saline with 0.2% Tween-20), then incubated overnight at 4°C with primary antibodies diluted in 5% BSA. The complete list of antibodies used for Western blotting is provided in Table 2. After washing, membranes were probed with appropriate HRP-conjugated secondary antibodies for 1 h at room temperature. Protein bands were visualized using enhanced chemiluminescence substrate (Clarity ECL substrate, 170-5060, Bio-Rad) and imaged with the ChemiDoc imaging system. Band intensities were quantified using Image Lab software (version 6.0.1, Bio-Rad). Non-phosphorylated proteins were normalized to stain-free total protein, whereas phosphorylated targets were expressed relative to the corresponding total protein.

**Table 2:** List of antibodies used for Western blotting (WB) and immunohistofluorescence (IF)

| Target proteins | Use | Dilution | Source, # product no. |
| --- | --- | --- | --- |
| Rabbit anti-Akt(pan) | WB | 1:3000 | Cell Signaling Technology, #4691 |
| Rabbit anti-ERK1/2 | WB | 1:3000 | Cell Signaling Technology, #9102 |
| Rabbit anti-GAP43 | WB | 1:3000 | Cell Signaling Technology, #8945 |
| Mouse anti-GFAP | WB | 1:3000 | Sigma Aldrich, #G3893 |
|  | IF | 1:150 |  |
| Goat anti-Iba1 | WB | 1:3000 | Abcam, #ab5076 |
| Mouse anti- IL-1β | WB | 1:1000 | Cell Signaling Technology, #12242 |
| Rabbit anti-IL-6 | WB | 1:1000 | Cell Signaling Technology, #12912 |
| Rabbit anti-JNK | WB | 1:3000 | Cell Signaling Technology, #9252 |
| Rabbit anti-laminin | IF | 1:750 | Sigma Aldrich, #L9393 |
| Rabbit anti-mTOR | WB | 1:3000 | Cell Signaling Technology, #2972 |
| Mouse anti-MyoD1 | WB | 1:1000 | Abcam, #Ab16148 |
| Rabbit anti-P-Akt(S473) | WB | 1:3000 | Cell Signaling Technology, #4060 |
| Rabbit anti-P-ERK1/2(T202/Y204) | WB | 1:3000 | Cell Signaling Technology, #4370 |
| Rabbit anti-P-Glucocorticoid receptor(S211) | WB | 1:3000 | Cell Signaling Technology, #4161 |
| Rabbit anti-P-JNK(T183/Y185) | WB | 1:3000 | Cell Signaling Technology, #4668 |
| Rabbit anti-P-mTOR(S2448) | WB | 1:3000 | Cell Signaling Technology, #2971 |
| Rabbit anti-P-TrkB(Y515) | WB | 1:3000 | GeneTex, #GTX00804 |
| Rabbit anti-NGFR | WB | 1:3000 | Fine Test, #FNab06091 |
| Mouse anti-PAX7 | WB | 1:3000 | Abcam, #Ab199010 |
| Rabbit anti-BDNF | WB | 1:3000 | Abcam, #Ab108319 |
| Rabbit anti-PSD95 | WB | 1:3000 | Cell Signaling Technology, #3450 |
| Rabbit anti-synaptophysin | WB | 1:3000 | ThermoFisher, #Epredia RB1461-P1 |
| Mouse anti-TNF-α | WB | 1:1000 | Cell Signaling Technology, #11948 |
| Mouse anti-TrkB-FL | WB | 1:3000 | BD Biosciences, #610102 |
| Rabbit anti-vimentin | WB | 1:1500 | Cell Signaling Technology, #5741 |
| HRP-conjugated secondary anti-rabbit | WB | 1:20000 | Jackson ImmunoResearch, #111-035-144 |
| HRP-conjugated secondary anti-mouse | WB | 1:20000 | Jackson ImmunoResearch, #115-035-166 |
| HRP-conjugated secondary anti-goat | WB | 1:10000 | Abcam, #ab6741 |
| Alexa Fluor 488 secondary goat anti-rabbit | IF | 1:500 | ThermoFisher, #A11008 |
| Alexa Fluor 594 secondary goat anti-rabbit | IF | 1:500 | ThermoFisher, #A21125 |
| CF Dye 568 secondary goat anti-mouse | IF | 1:500 | VWR Biotium, #20102 |

### 2.11. Skeletal muscle and brain sample sectioning for histological analyses

Quadriceps muscle samples were mounted in tragacanth, frozen in precooled isopentane, and then stored at −80°C. Serial quadriceps cross-sections, 10 µm in thickness, were cut at −20°C using a cryostat and mounted on lysine-coated Superfrost slides. Brain samples were embedded in paraffin, cut into 5-µm-thick coronal sections through the hippocampus using an HM325 microtome (MicroM, Brignais, France), and mounted on lysine-coated Superfrost slides.

### 2.12. Histological and immunofluorescence analyses

#### 2.12.1. Skeletal muscle

Quadriceps cross-sections were brought to room temperature, rehydrated in phosphate-buffered saline (PBS; Sigma, 4417), and permeabilized in PBS containing 0.1% Triton X-100 for 15 min. Sections were blocked in PBS containing 10% normal goat serum for 30 min at room temperature and incubated for 1 h with an anti-laminin primary antibody diluted in blocking buffer. After three PBS washes, sections were incubated for 1 h with the appropriate secondary antibody (see Table 2 for antibody details). Sections were then washed three times in PBS and coverslipped using Fluoro-gel mounting medium containing DAPI (FP-DT094A, Interchim). Slides were imaged using a fluorescence microscope (Axio Imager, Zeiss, Oberkochen, Germany). Myofiber cross-sectional area and the percentage of centrally nucleated fibers were quantified using the MuscleJ2 plugin (Danckaert *et al*., 2023) and manual counting, respectively. Fiber-level measurements were averaged within each animal before statistical analysis. At least 500 myofibers per animal were evaluated to determine the percentage of centrally nucleated fibers.

Muscle collagen content was assessed by second harmonic generation (SHG) microscopy using a Nikon AX R MP multiphoton microscope equipped with a femtosecond pulsed laser and a ×40 objective. Samples were excited at 920 nm, and the SHG signal was collected in epi-detection mode within a 440– 480 nm detection window. Five non-overlapping fields were acquired from anatomically matched quadriceps cross-sections for each animal using identical acquisition settings. Images were acquired with NIS-Elements AR software (Nikon). The area occupied by collagen-associated SHG signal was segmented using Li’s automatic thresholding method, applied uniformly without manual adjustment, and expressed relative to the total tissue area analyzed. Measurements from the five fields were averaged to obtain a single value per animal, which was considered the statistical unit. Image acquisition and quantification were performed blinded to group allocation.

#### 2.12.2. Brain

For GFAP immunofluorescence, paraffin-embedded brain sections containing the hippocampus were first deparaffinized using standard xylene and graded ethanol baths, then rinsed in water. Heat-induced epitope retrieval was performed by immersing the slides in preheated 10 mM sodium citrate buffer (pH 6.0) at 95°C for 15 min; sections were then allowed to cool to room temperature in the same buffer for 20 min. After washing in PBS, sections were blocked with 3% bovine serum albumin (BSA) in PBS. Sections were incubated overnight at 4°C with the primary antibody. After three washes in PBS, sections were incubated at room temperature for 1 h with the appropriate secondary antibody. After final washes, sections were mounted with a DAPI-containing mounting medium (FP-DT094A, Interchim). Slides were imaged using a fluorescence microscope (Axio Imager, Zeiss, Oberkochen, Germany). Analyses were restricted to the hippocampus, including the dentate gyrus (DG), CA1, and CA3 subregions. Quantification was performed using a custom image-analysis macro developed in-house, based on Triangle threshold method, available at [https://github.com/JulienWIRTZ/Astrocyte_GFAP_Skeleton] and archived at [10.5281/zenodo.23057922].

For Nissl staining, coronal brain sections were deparaffinized in xylene, rehydrated through graded ethanol solutions, and rinsed in distilled water. Sections were stained with 0.1% cresyl violet (5042-10G, Sigma Aldrich) solution for 7 min, differentiated in 95% ethanol containing 1% acetic acid for 24 s, dehydrated through graded ethanol, cleared in xylene and mounted with a permanent mounting medium (Eukitt, 92537). Neuronal profiles were quantified semi-automatically in the granule cell layer of the dentate gyrus and in the pyramidal cell layers of CA1 and CA3. For each animal, two anatomically matched sections were imaged under identical acquisition settings using a ×40 objective. Regions of interest were manually delineated in ImageJ software. Neuronal profiles were identified by the presence of a large, round or oval, lightly stained nucleus, whereas profiles containing small, intensely stained nuclei with glial-like morphology were excluded. The number of neuronal profiles was normalized to the analyzed area. Values obtained from the two sections were averaged, and each animal was considered the statistical unit. Because counts were obtained from single-plane images rather than by design-based stereology, these values are reported as neuronal profile density and do not provide an unbiased estimate of total neuronal number.

### 2.13. Statistical analysis

Data are presented as individual values with mean ± SD, with each animal considered an independent biological replicate. Normality was assessed using the Shapiro-Wilk test together with graphical inspection of data distributions, and equality of variances was evaluated using an F-test. Comparisons between two independent groups were performed using two-tailed unpaired Student’s t-tests when parametric assumptions were met, Welch’s t-tests when variances were unequal, or Mann-Whitney U tests for non-normally distributed data. Longitudinal and distribution data were analyzed using two-way repeated-measures ANOVA with experimental group as the between-subject factor and, as the within-subject factor, time for body mass, cross-sectional area interval for fiber size distributions, and 1-min interval for forced-swim immobility. The Geisser-Greenhouse correction was applied when appropriate, followed by Šidák-adjusted multiple comparisons when a significant interaction was detected. The proportions of samples with detectable serum IL-6 concentrations were compared using a two-sided Fisher’s exact test. Associations between variables were assessed across all animals pooled, using Pearson’s correlation for normally distributed data and Spearman’s rank correlation otherwise. No correction for multiple comparisons was applied across the independent endpoints examined, and individual comparisons are therefore interpreted in the context of the overall pattern rather than in isolation. Exact p-values are reported whenever possible and statistical significance was set at p < 0.05. Analyses were performed using GraphPad Prism version 9.5.0 (GraphPad Software, San Diego, CA, USA).

### 2.14. Declaration of generative AI use

During the preparation of this manuscript, the authors used Claude (Anthropic) to obtain structured editorial feedback on successive drafts, including suggestions for the organisation and wording of the Introduction and Discussion, cross-checking of the text against the figures, and verification of bibliographic references against PubMed records. The tool was not used to generate, analyse or interpret scientific data. All study design, data analysis, scientific interpretation and conclusions are the authors’ own. The authors reviewed and edited all AI-assisted output and take full responsibility for the content of the article.

## 3. Results

### 3.1. Physiological effects of repeated NMES

Body mass remained comparable between groups at each of the three time points examined (Fig. 1B), and body mass-normalized quadriceps mass did not differ at the end of the protocol (Fig. 1C). In the behavioral cohort, spontaneous locomotor activity, quantified as the number of grid crossings, was likewise unchanged (Fig. 1D). By contrast, NMES-treated animals recovered more rapidly from anesthesia than SHAM controls, as shown by the shorter post-anesthetic recovery time (Fig. 1E). Since both groups received an identical anesthetic protocol at each session, this difference is attributable to the evoked contraction rather than to anesthetic exposure itself.

### 3.2. Muscle remodeling and collagen deposition accompanied by low-grade systemic inflammation

To determine whether repeated NMES altered the structural integrity of the stimulated muscle, circulating creatine kinase (CK) activity and quadriceps histology were examined. CK activity was elevated immediately after the final NMES session, consistent with transient sarcolemmal perturbation (Fig. 2A). At 24 h, CK activity remained higher in NMES-treated animals but no longer reached statistical significance (p = 0.0528; Fig. 2A). Repeated NMES was also associated with an increased collagen-associated SHG signal in the stimulated quadriceps compared with SHAM controls (Fig. 2B). This alteration was not accompanied by an increase in centrally nucleated fibers, the proportion of which remained comparable between groups (Fig. 2C), nor by muscle hypertrophy, the distribution of myofiber cross-sectional areas being similar between groups (Fig. 2D). In addition, the protein expression of PAX7 and MyoD1, markers associated with satellite-cell maintenance and myogenic differentiation, respectively, was unchanged (Fig. 2E-F). These findings indicate that the increased collagen signal occurred without changes in myofiber size or in the expression of the examined myogenic markers at the time of tissue collection. The response to repeated NMES was not restricted to the stimulated muscle. Serum IL-6 was below the assay detection limit in all SHAM animals but detectable in all NMES-treated animals (0/4 vs. 4/4 detectable samples; two-sided Fisher’s exact test, p = 0.0286). Serum TNF-α concentrations were also increased, whereas serum IL-1β concentrations remained unchanged (Fig. 2G-I).

**Figure 2.**
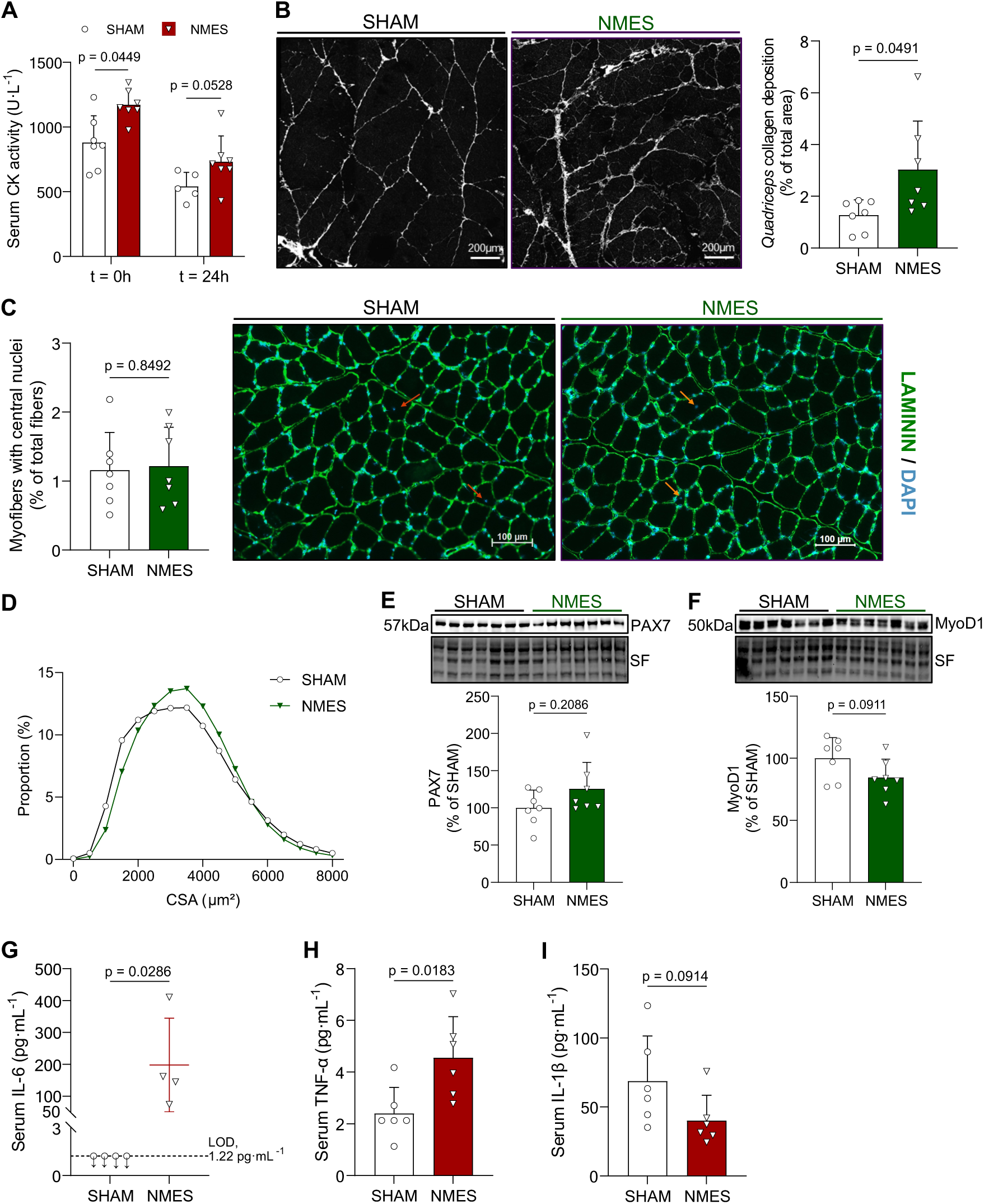
Repeated NMES induces muscle perturbation and a low-grade systemic inflammatory response. (A) Serum creatine kinase activity measured immediately after the final session (0 h) and 24 h later. (B) Representative second-harmonic generation images of quadriceps fibrillar collagen and quantification of collagen-positive area. Scale bar, 200 µm. (C) Percentage of myofibers containing centrally located nuclei and representative laminin/DAPI-stained quadriceps cross-sections. Orange arrows indicate centrally nucleated fibers. Scale bar, 100 µm. (D) Cross-sectional area (CSA) distribution of muscle fibers expressed as relative frequency (%). (E-F) Representative Western blots and quantification of PAX7 and MyoD1 expression. Stain-free total protein was used as the loading control. (G-H-I) Serum IL-6, TNF-α and IL-1β concentrations; in (G), the dashed line indicates the assay limit of detection. Data are presented as individual values with mean ± SD. IL-6 detectability was compared using a two-sided Fisher’s exact test. Other comparisons were performed using the appropriate two-tailed unpaired test.

### 3.3. Altered stress-related behavior and enhanced glucocorticoid signaling

The behavioral consequences of short-term repeated NMES were assessed using tasks evaluating recognition memory and stress-related behavioral responses. During the novel object recognition test, the discrimination index did not differ between NMES-treated and SHAM animals in either the short-term or the long-term retention session (Fig. 3A-B). Total object exploration time was also comparable at every phase of the task, indicating similar engagement with the objects (Appendice Fig. 1). In contrast, repeated NMES altered responses in behavioral paradigms sensitive to repetitive digging and coping strategies. NMES-treated animals buried more marbles than SHAM controls in the marble-burying test (Fig. 3C) and displayed a longer duration of immobility in the forced swim test (Fig. 3D). Analysis of immobility across successive one-min intervals revealed a significant time-by-group interaction, indicating that the temporal pattern of immobility differed between NMES-treated and SHAM animals [F(5,60) = 2.763, p = 0.0260]. Šídák-adjusted post hoc comparisons showed greater immobility in NMES-treated animals during the 2-3, 4-5 and 5-6 min intervals (Fig. 3E). As spontaneous locomotor activity was unchanged between groups, these behavioral differences were unlikely to be explained by a generalized reduction in basal movement. Repeated NMES was also associated with increased glucocorticoid signaling. Circulating corticosterone concentrations were higher in NMES-treated animals 24 h after the final stimulation session (Fig. 3F), and hippocampal glucocorticoid receptor phosphorylation at Ser211 was increased in the NMES group (Fig. 3G). Furthermore, circulating corticosterone concentrations were positively correlated with hippocampal glucocorticoid receptor phosphorylation across all animals for which both measurements were available (n = 12; r = 0.7109, p = 0.0095, Fig. 3H).

**Figure 3.**
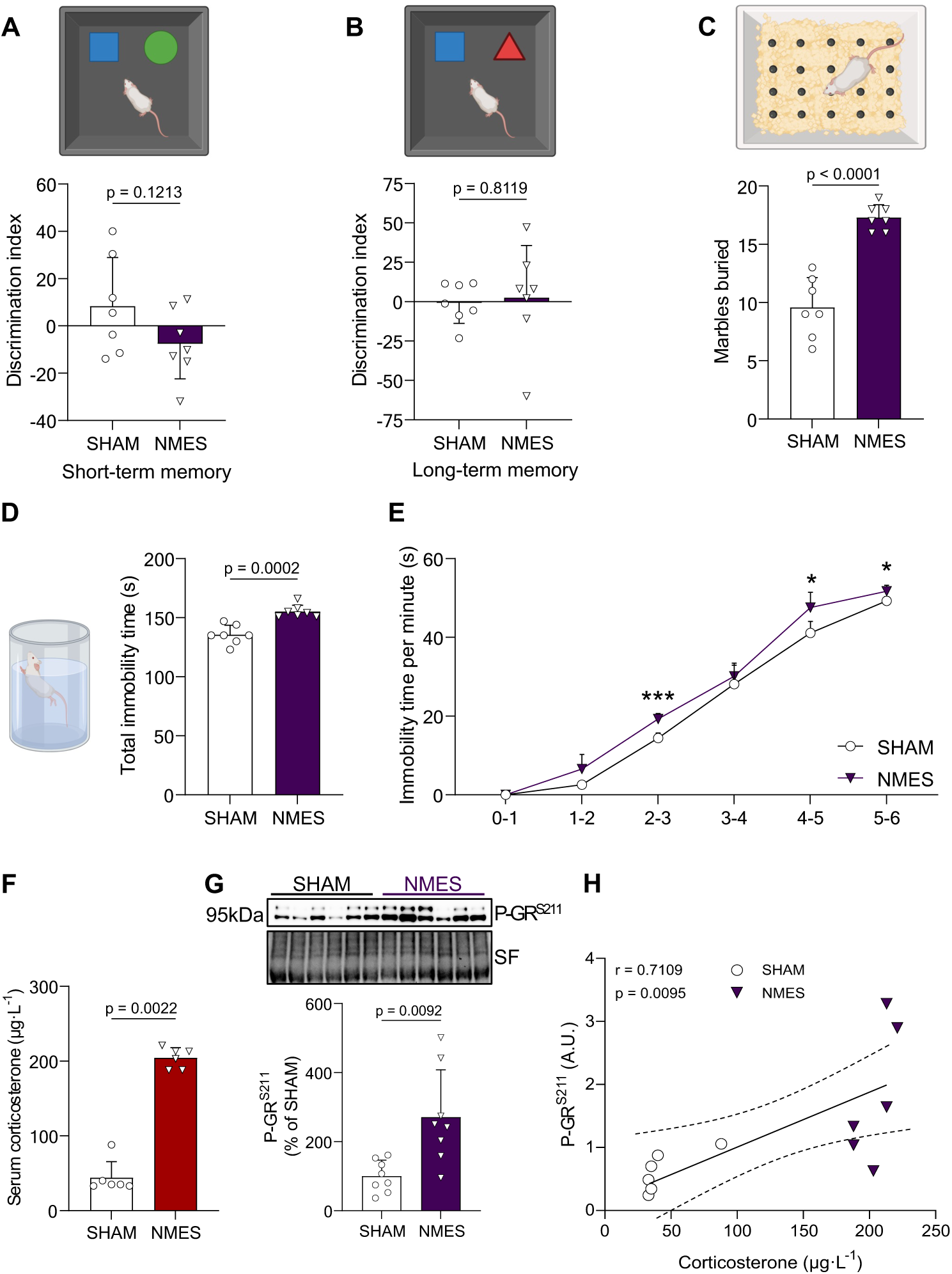
Repeated NMES selectively alters behavioral outcomes and enhances glucocorticoid signaling. (A-B) Novel object recognition performance assessed by the discrimination index during the short- and long-term memory tests. (C) Number of marbles buried during the marble-burying test. (D) Total immobility time during the forced-swim test. (E) Immobility time across successive 1-min intervals of the forced-swim test. (F) Serum corticosterone concentration measured 24 h after the final NMES session. (G) Representative Western blots and quantification of hippocampal glucocorticoid receptor phosphorylation at Ser211. Stain-free total protein was used as the loading control. (H) Association between serum corticosterone and hippocampal P-GR^Ser211^ expression, computed across all animals from both groups. The solid line represents the linear regression and dashed lines the 95% confidence interval. Data are presented as individual values with mean ± SD. Asterisks in (E) indicate statistically significant differences: * *p < 0.05, *** p < 0.001.* Minute-by-minute immobility was analyzed using two-way repeated-measures ANOVA with Geisser-Greenhouse correction and Šidák-adjusted comparisons. The association in (H) was assessed using Pearson’s correlation. Other comparisons were performed using the appropriate two-tailed unpaired test.

### 3.4. Widespread glial and inflammatory response in the hippocampus

Glial and inflammatory responses were then assessed in the hippocampus. Repeated NMES was associated with a 33% higher mean GFAP protein expression by Western blot, although this difference did not reach statistical significance (p = 0.0767), and significantly increased vimentin expression (Fig. 4A-B). Hippocampal Iba1 expression was also elevated in NMES-treated animals (Fig. 4C). The astrocytic response was more clearly detected by immunofluorescence, which revealed a significantly greater GFAP-positive area in the dentate gyrus (DG), CA1, and CA3 regions of NMES-treated animals (Fig. 4D). The response was therefore distributed across hippocampal subfields. In parallel, hippocampal *Il1b*, *Il6*, and *Tnf* mRNA levels were increased following repeated NMES (Fig. 4E-G). These transcriptional changes were accompanied by higher hippocampal IL-1β and IL-6 protein expression in the NMES group (Fig. 4H), while both precursor and mature TNF-α protein forms were likewise increased (Fig. 4H). Although the cellular sources of these cytokines were not determined, these molecular changes, together with the increased expression of GFAP, vimentin, and Iba1, support a coordinated hippocampal glial and inflammatory response to repeated NMES.

**Figure 4.**
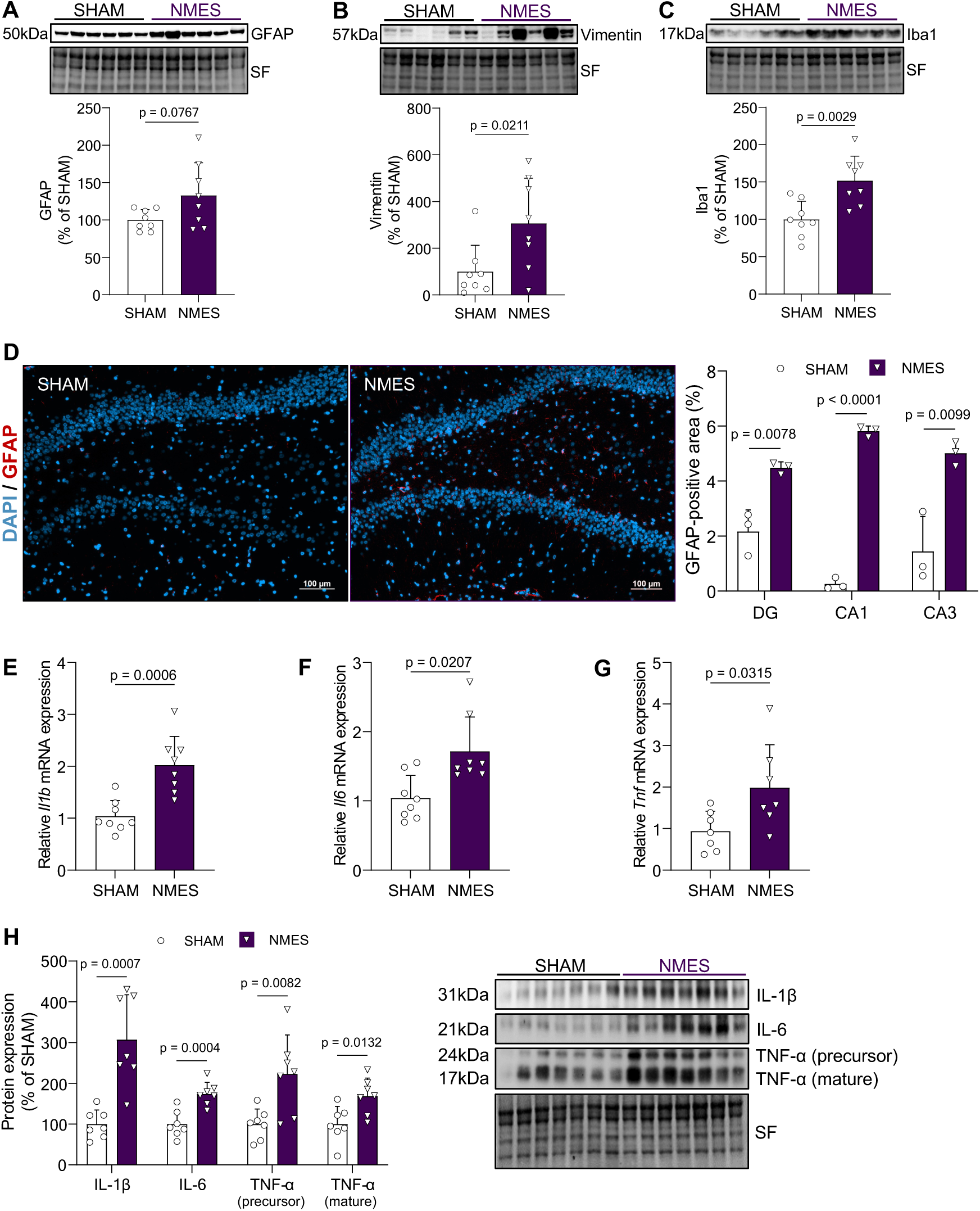
Repeated NMES induces hippocampal glial and inflammatory responses. (A-C) Representative Western blots and quantification of hippocampal GFAP, vimentin, and Iba1 expression. Stain-free total protein was used as the loading control. (D) Representative GFAP/DAPI-stained hippocampal sections, together with quantification of GFAP-positive area in the dentate gyrus (DG), cornu ammonis 1 (CA1), and CA3. Scale bars, 100 µm. (E-G) Relative hippocampal *Il1b*, *Il6*, and *Tnf* mRNA expression. (H) Representative Western blots and grouped quantification of IL-1β, IL-6, and precursor and mature TNF-α expression. Stain-free total protein was used as the loading control. Data are presented as individual values with mean ± SD. Comparisons were performed using the appropriate two-tailed unpaired test.

### 3.5. Hippocampal BDNF signaling shifts toward the proBDNF-p75^NTR^-JNK axis

The effect of repeated NMES on hippocampal BDNF signaling was subsequently examined. *Bdnf* mRNA expression did not differ between NMES-treated and SHAM animals (Fig. 5A). At the protein level, however, the precursor and mature forms of BDNF were differentially regulated. Mature BDNF expression was reduced, whereas proBDNF expression was increased in the hippocampus of NMES-treated animals (Fig. 5B-C). As a result, the proBDNF-mBDNF ratio was increased following repeated NMES (Fig. 5D). This shift in BDNF protein forms was accompanied by changes in receptor-associated signaling. TrkB phosphorylation at Tyr515 was reduced in the NMES group (Fig. 5E), together with a decrease in ERK1/2 phosphorylation (Fig. 5F). In parallel, p75^NTR^ protein expression was increased following repeated NMES (Fig. 5G), accompanied by elevated JNK phosphorylation (Fig. 5H). This rebalancing did not extend to all downstream pathways, since neither Akt nor mTOR phosphorylation differed between groups (Fig. 5I-J). Thus, repeated NMES was associated with a selective reduction in mBDNF-TrkB-ERK-related signaling together with increased proBDNF-p75NTR-JNK-related signaling, while Akt-mTOR signaling remained unchanged.

**Figure 5.**
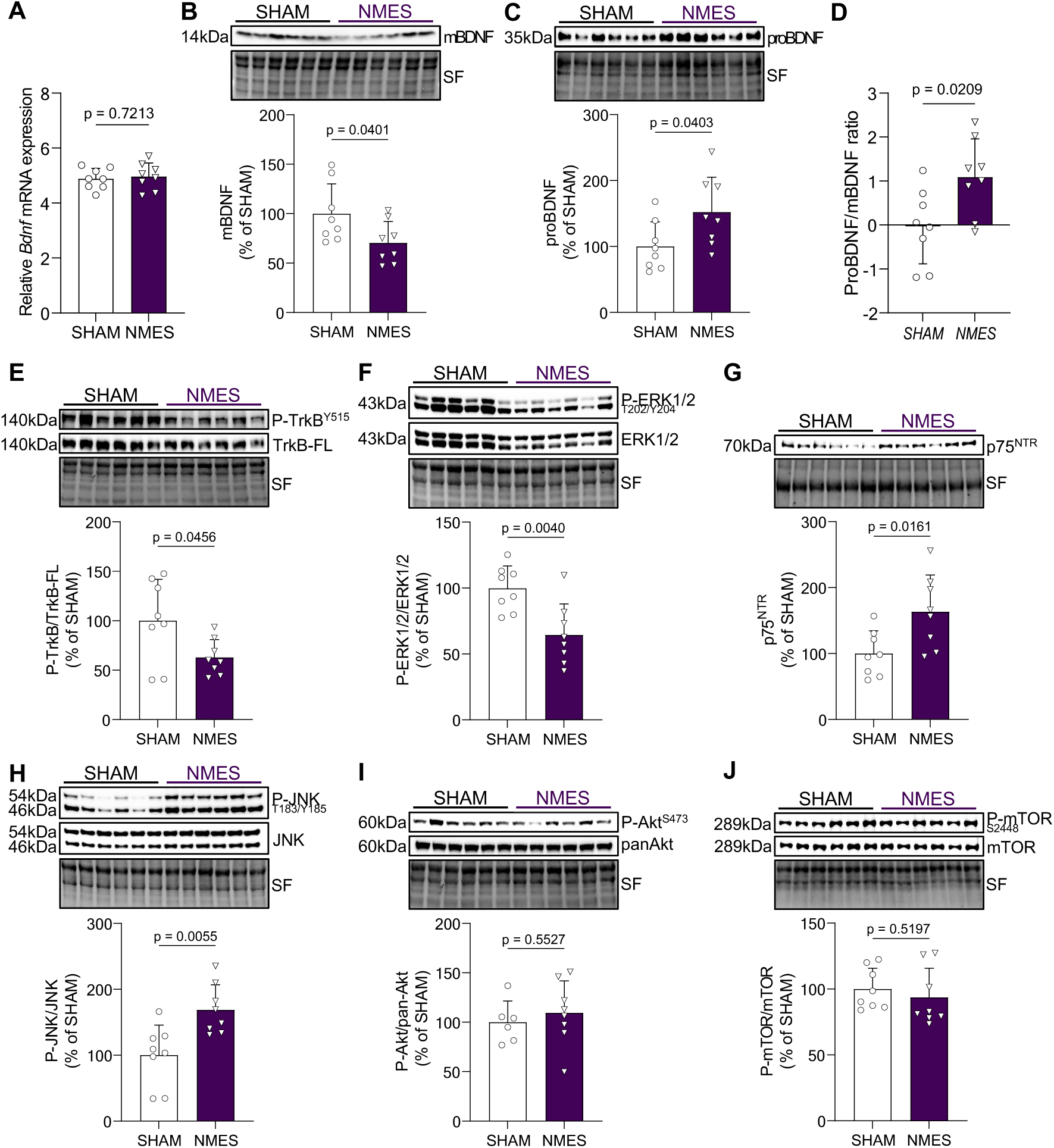
Repeated NMES shifts hippocampal BDNF signaling toward the proBDNF-p75^NTR^-JNK axis. (A) Relative hippocampal *Bdnf* mRNA expression. (B-C) Representative Western blots and quantification of mature BDNF and proBDNF expression. (D) Balance between the two BDNF forms, expressed as the log2 ratio of proBDNF to mature BDNF, each normalized to the SHAM mean. Positive values indicate a shift toward proBDNF, negative values a shift toward mature BDNF. (E) TrkB phosphorylation at Tyr515 normalized to full-length TrkB. (F) ERK1/2 phosphorylation at Thr202 and Tyr204 normalized to total ERK1/2. (G) p75^NTR^ expression. (H) JNK phosphorylation at Thr183 and Tyr185 normalized to total JNK. (I) Akt phosphorylation at Ser473 normalized to pan-Akt. (J) mTOR phosphorylation at Ser2448 normalized to total mTOR. Representative Western blots are shown above their corresponding quantifications, and stain-free total protein was used as the loading control where appropriate. Data are presented as individual values with mean ± SD and were analyzed using the appropriate two-tailed unpaired test.

### 3.6. Reduced plasticity-related proteins without altered neuronal profile density

To determine whether the changes in BDNF signaling were accompanied by alterations in proteins associated with axonal and synaptic plasticity, synaptophysin (SYP), GAP43, and PSD95 were quantified in the hippocampus. Synaptophysin, a presynaptic vesicle-associated protein, was reduced in NMES-treated animals compared with SHAM controls (Fig. 6A). GAP43 expression was likewise decreased following repeated NMES, together with a reduction in the postsynaptic scaffold protein PSD95 (Fig. 6A). This concomitant reduction indicates that the molecular changes involved presynaptic, axonal plasticity-associated, and postsynaptic proteins. Nissl staining was then used to assess neuronal profile density. The density of Nissl-positive neuronal profiles remained unchanged in the dentate gyrus, CA1, and CA3 regions (Fig. 6B). The reduction in plasticity-related proteins therefore occurred without detectable neuronal loss.

**Figure 6.**
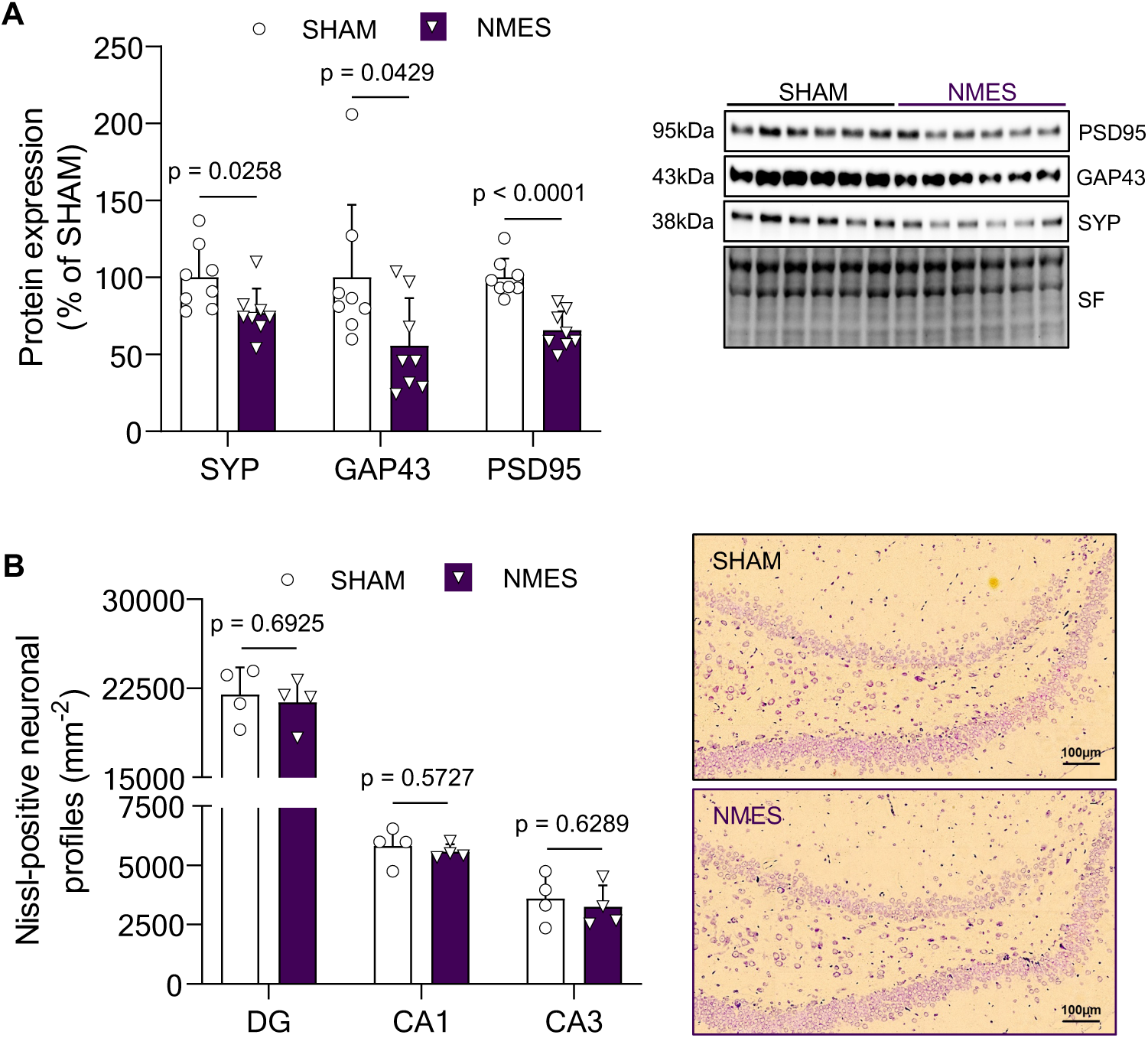
Repeated NMES reduces hippocampal synaptic proteins without a detectable reduction in neuronal profile density. (A) Representative Western blots and grouped quantification of synaptophysin (SYP), GAP43, and PSD95 expression in the hippocampus. Stain-free total protein was used as the loading control. (B) Quantification of Nissl-positive neuronal profiles in the dentate gyrus (DG), cornu ammonis 1 (CA1), and CA3, with representative Nissl-stained sections from SHAM and NMES animals. Values are profile densities obtained from single-plane images and are not stereological estimates of neuronal number. Scale bar, 100 µm. Data are presented as individual values with mean ± SD. Comparisons were performed using the appropriate two-tailed unpaired test.

## 4. Discussion

Contrary to our hypothesis that daily stimulation would reproduce the beneficial muscle-to-brain signaling reported after an acute NMES protocol (Chaney *et al*., 2024), seven consecutive days of quadriceps NMES induced a coordinated peripheral and hippocampal response that diverged from the trophic adaptations typically associated with voluntary exercise. Repeated NMES sessions reduced mature BDNF-TrkB-ERK signaling, drove a reciprocal increase in proBDNF-p75^NTR^-JNK signaling, elevated circulating corticosterone, triggered hippocampal glial and inflammatory activation, and produced a shift in stress-related behavior. Critically, these central changes occurred without a detectable reduction in neuronal profile density and alongside only moderate skeletal muscle remodeling, with no changes in muscle mass, myofiber cross-sectional area, or the myogenic markers PAX7 and MyoD1 expression. These findings indicate that repeated NMES does not simply prolong the acute response, but engages a distinct, maladaptive mode of muscle-to-brain signaling, despite a limited footprint in the skeletal muscle itself.

At the peripheral level, repeated NMES produced a muscle response consistent with peripheral perturbation rather than overt structural injury. Circulating creatine kinase increased immediately after the seventh and final session of NMES and remained elevated at 24 h but no longer significantly different from the SHAM group. This profile is compatible with transient sarcolemmal stress and possibly incomplete recovery (Nosaka *et al*., 2011). The increased collagen-associated SHG signal should be interpreted as reflecting early extracellular-matrix remodeling rather than established fibrosis, given the absence of myofiber damage (unchanged fiber size, centrally nucleated fiber proportion, PAX7/MyoD1 expression). This pattern indicates repeated peripheral muscle perturbation without overt myofiber degeneration-regeneration cycling and stands in contrast to the more pronounced structural muscle damage observed in the soleus, and to a lesser extent the gastrocnemius, after lumbar-nerve stimulation (Wirtz *et al*., 2025). Twenty-four hours after the last session, circulating TNF-α was elevated, and IL-6 was detectable in NMES animals while remaining below the detection limit in SHAM animals. Contracting muscle is a major source of circulating IL-6 (Pedersen & Febbraio, 2008), making the stimulated quadriceps a plausible contributor, although other sources cannot be excluded. Their pattern is more informative than their origin. During voluntary exercise, the IL-6 response is transient, resolving within hours, and occurs without a rise in TNF-α (Pedersen & Febbraio, 2008). Their joint elevation a full day after stimulation therefore points to an inflammatory state rather than the physiological myokine signature of exercise. Circulating IL-1β, by contrast, was unchanged. The peripheral profile is thus best described as a low-grade inflammatory state, consistent in magnitude with the moderate muscle perturbation documented above.

As NMES and SHAM animals underwent identical anesthesia, handling and timing, evoked contraction was the only difference between groups. Repeated evoked contraction, even without overt muscle injury, is therefore sufficient to initiate a peripheral signal capable of reaching the hippocampus. A sustained glucocorticoid response was among the most prominent systemic consequences of repeated NMES. Circulating corticosterone and hippocampal glucocorticoid receptor phosphorylation at Ser211 were both elevated 24 h after the final session and positively correlated across animals, although this correlation reflects the group difference (NMES vs SHAM) rather than a dose-dependent relationship within either group. Obtained under terminal anesthesia, and with animals anesthetized throughout each stimulation session, this persisting elevation is unlikely to reflect a conscious or psychological stress component, and more plausibly reflects the reactive arm of the neuroendocrine stress response, driven directly by sensory relays and humoral signals from the repeatedly contracted muscle (Herman *et al*., 2016). Habituation of the glucocorticoid response to repeated homotypic stress is only partial (Grissom & Bhatnagar, 2009), consistent with an elevation still detectable a day after the seventh daily session. This glucocorticoid signature was paralleled by a selective behavioral shift. Repeated NMES increased marble burying and forced-swim immobility without altering spontaneous locomotion, so these changes are unlikely to reflect reduced general mobility. Recognition memory was not detectably affected. The forced-swim finding parallels the well-established sensitivity of this behavior to sustained corticosterone elevation, which increases immobility in male rats without altering general activity (Gregus *et al*., 2005), while a comparable co-occurrence of increased forced-swim immobility and altered marble-related behavior under chronic corticosterone elevation has also been reported, in female rats (Conrad *et al*., 2024). Because behavioral and molecular endpoints were obtained from separate animal cohorts, the behavioral change cannot be linked to the hippocampal signature within individual animals. The behavioral panel, moreover, does not cover the dimensions required to establish an anxiety-or depression-like syndrome, so the effect is best described as a selective shift in affect-related behavior. Together, these data identify glucocorticoid receptor activation as a candidate mediator of the central changes described below.

Indeed, the glial response was consistent with the glucocorticoid signature. Increased Iba1 and vimentin protein expression, together with an expanded GFAP-positive area across hippocampal subfields, indicate coordinated microglial and astrocytic remodeling; whole-hippocampus GFAP protein expression showed only a non-significant trend, consistent with a regionally distributed rather than a globally uniform astrocytic response. The intensity of this reaction is notable, since vimentin is upregulated together with GFAP as part of the intermediate-filament signature of reactive astrogliosis (Pekny & Pekna, 2014), while the microglial component is consistent with the sensitization toward a pro-inflammatory phenotype produced by sustained glucocorticoid signaling (Frank *et al*., 2012). Because chronic psychological or purely glucocorticoid-driven stress typically produces astrocyte atrophy rather than overt reactivity (Tynan *et al*., 2013), this reactive profile points to an additional, cytokine-driven component rather than a glucocorticoid-only origin. The repeatedly stimulated muscle may drive this component through at least two non-exclusive routes. The first is humoral, since circulating IL-6 and TNF-α, both elevated after repeated NMES, are plausible candidates for reaching the hippocampus and engaging resident glia (Novakovic *et al*., 2023). The second, more speculative, is metabolic, since excessive muscle demand may impair the physiological diversion of kynurenine toward kynurenic acid (Agudelo *et al*., 2014) and instead raise central kynurenine availability, favoring its microglial conversion to the astrogliotoxic metabolite quinolinic acid (Guillemin, 2012; Da Rocha *et al*., 2025). Neither route, however, accounts for the full hippocampal profile, since IL-1β, which remained unchanged in the circulation as noted above, was nonetheless induced in the hippocampus at both transcript and protein level. This dissociation points to local synthesis rather than passive accumulation of a circulating pool (Dantzer *et al*., 2008), and the parallel induction of Il6 and Tnf transcripts suggests that the same holds for the other two cytokines. Although the cellular source of these cytokines cannot be determined here, this centrally produced component is plausibly microglial. Rather than replacing the peripheral routes, such local production would complement them, and since activated microglia are themselves established inducers of neurotoxic reactive astrocytes (Liddelow *et al*., 2017), peripheral and local signals may converge on a single microglia-to-astrocyte cascade, a possibility that remains untested here but is directly relevant to muscle-to-brain signaling and warrants dedicated investigation.

Hippocampal neuroplasticity was the most tightly coordinated readout of this maladaptive cascade, spanning BDNF signaling itself and the synaptic proteins it supports. Mature BDNF, TrkB and ERK1/2 phosphorylation were reduced while proBDNF, p75^NTR^ and JNK phosphorylation increased, shifting the proBDNF/mBDNF balance toward a profile favoring synaptic weakening over potentiation (Woo *et al*., 2005; Li *et al*., 2019). The unchanged Bdnf transcript points to a post-transcriptional origin for this shift, and two non-exclusive mechanisms can account for it. Sustained glucocorticoid receptor activation is a well-established driver, increasing proBDNF synthesis through an endoplasmic-reticulum-stress-dependent mechanism (Liu *et al*., 2020) and reducing the proteolytic conversion of proBDNF to mature BDNF through a fall in hippocampal tissue plasminogen activator (Yeh *et al*., 2012). The hippocampal cytokine elevation described above offers a second route to the same endpoint. IL-1β impairs the retrograde trafficking of BDNF signaling endosomes and the downstream phosphorylation they carry, producing a state of neurotrophic resistance without altering BDNF transcription (Carlos *et al*., 2017). Consistently, blocking central IL-1 receptor signaling prevents the reduction in mature BDNF and TrkB-ERK activation after a peripheral immune challenge, at least in aged animals (Cortese *et al*., 2011). This rebalancing was selective, as Akt and mTOR phosphorylation were unchanged. Reduced hippocampal ERK1/2 signaling has likewise been reported under uncontrollable and chronic mild stress (Dwivedi & Zhang, 2016; Molteni *et al*., 2016), whereas more severe paradigms such as chronic restraint suppress Akt and mTOR as well (Seo *et al*., 2017). It was mirrored at the structural level by a concurrent reduction in synaptophysin, GAP43 and PSD95 across presynaptic, axonal-growth and postsynaptic compartments, without a detectable change in neuronal profile density. This pattern is consistent with p75^NTR^ signaling driving synapse elimination and dendritic regression independently of neuronal death (Zagrebelsky *et al*., 2005; Egashira *et al*., 2010). However, reduced protein abundance does not necessarily equate to reduced synaptic function, and the single 24-h time point and the absence of direct functional measures cannot exclude a reversible course for this synaptopathy, since stress-induced structural plasticity changes of this kind are generally transient and reversible once the stressor resolves (Lucassen *et al*., 2006).

Several limitations should be considered when interpreting these findings. As noted above, the single 24-h time point leaves the reversibility of the behavioral and synaptic changes unresolved and precludes reconstructing their temporal sequence. The peripheral signal reaching the hippocampus may also not originate exclusively from the myofiber, since NMES recruits motor and sensory axons and can elicit its own metabolic, vascular and endocrine responses (Enoka *et al*., 2020). Within the core BDNF signaling panel, several comparisons cleared significance only narrowly (mature BDNF, p = 0.0401; proBDNF, p = 0.0403; TrkB phosphorylation, p = 0.0456; GAP43, p = 0.0429). Taken individually these comparisons remain exploratory, and confidence in the overall direction of the effect rests instead on their convergence with the more robustly significant changes in JNK and p75^NTR^ signaling, glial and cytokine markers, and behavior, all pointing toward the same maladaptive pattern. The associative design is a further limitation, as no intervention such as adrenalectomy or cytokine-signaling blockade was used to establish a necessary or sufficient mediator. Finally, the use of young male rats only limits the generalization of these findings, since sex and estrogens shape both the central stress response (Hodes *et al*., 2024) and the muscle-level inflammatory response to injury itself (Jomard & Gondin, 2023).

## 5. Conclusion

The present findings are consistent with the emerging view, synthesized at the muscle level in the context of overtraining syndrome by Shorter et al.(2026), that non-physiological or chronically excessive muscle stress engages a maladaptive state involving endocrine dysregulation and systemic inflammation. Our findings extend this state beyond the muscle itself, to the hippocampus and ultimately to affect-related behavior. This central phenotype arose despite comparatively modest muscle remodeling. It contrasts with the more severe lumbar-nerve stimulation protocol described above, in which pronounced muscle damage engaged glucocorticoid receptor signaling alone, without the systemic corticosterone rise or neuroinflammatory response documented here (Wirtz *et al*., 2025). This comparison suggests that the qualitative nature of the peripheral signal, rather than the extent of muscle damage, shapes the hippocampal outcome. That acute NMES instead increased hippocampal mature BDNF, as noted at the outset (Chaney *et al*., 2024), indicates that this maladaptive state is not an inevitable consequence of electrically evoked contraction. The per-session parameters were essentially those used here, differing only in stimulation frequency (40 Hz in this protocol versus 100 Hz in the acute one), with both regimens eliciting sustained tetanic contractions, so the divergent outcome is unlikely to stem from the stimulation parameters themselves and more plausibly reflects daily repetition without adequate recovery. Beyond session number and recovery, however, NMES offers a valuable approach for dissecting this beneficial-to-maladaptive switch, since current intensity, duration, frequency and target muscle (Gorgey *et al*., 2009; Behringer *et al*., 2016) are all fully tunable, making it uniquely suited to isolate the secretory programs that drive it. Until this switch is better understood, notably through modern secretome profiling (Ng & Handschin, 2026), caution is warranted before recommending repeated NMES as an exercise substitute for populations unable to exercise voluntarily, particularly older individuals, in whom muscle recovery from damaging exercise is delayed, prolonged and less efficient (Li *et al*., 2024).

## CrediT Authorship contribution statement

Julien Wirtz: conceptualization, data curation, formal analysis, investigation, methodology, writing – original draft, writing – review and editing. Yuan Wang: data curation, formal analysis, writing – review and editing. Chaney Rémi: data curation, methodology, investigation, writing – review and editing. Léa Wirtz: data curation, formal analysis, writing – review and editing. Quentin Drezet: data curation, writing – review and editing. Charalambos Papaxanthis: writing – review and editing. Aurore Quirié: data curation, formal analysis, writing – review and editing. Alexandre Méloux: data curation, writing – review and editing. Stéphanie Lemaire: data curation, formal analysis, investigation, writing – review and editing. Marina Cefis: conceptualization, data curation, methodology, formal analysis, investigation, methodology, supervision, writing – original draft, writing – review and editing. Anne Prigent-Tessier: investigation, supervision, writing – review and editing. Philippe Garnier: project administration, conceptualization, data curation, formal analysis, investigation, methodology, supervision, writing – original draft, writing – review and editing.

## Declaration of competing interest

The authors declare that they have no known competing financial interests or personal relationships that could have appeared to influence the work reported in this paper.

## Funding

This work was supported by the Region Bourgogne Franche-Comté, Envergure project MIND-STIM.

## Data availability

The image-analysis code used in this study is openly available at https://github.com/JulienWIRTZ/Astrocyte_GFAP_Skeleton and archived at DOI: 10.5281/zenodo.23057922. The data that support the findings of this study are available from the corresponding author upon reasonable request.

## Acknowledgments

We would like to thank the ImaFlow platform attached to the US58 BioSanD of the Université Bourgogne Europe for their help with immunolabeling experiments and the UMR1098 RIGHT—TAI-IT of the Université Marie & Louis Pasteur for multiplex analysis of circulating cytokines.

### Appendices

**Appendice 1.**
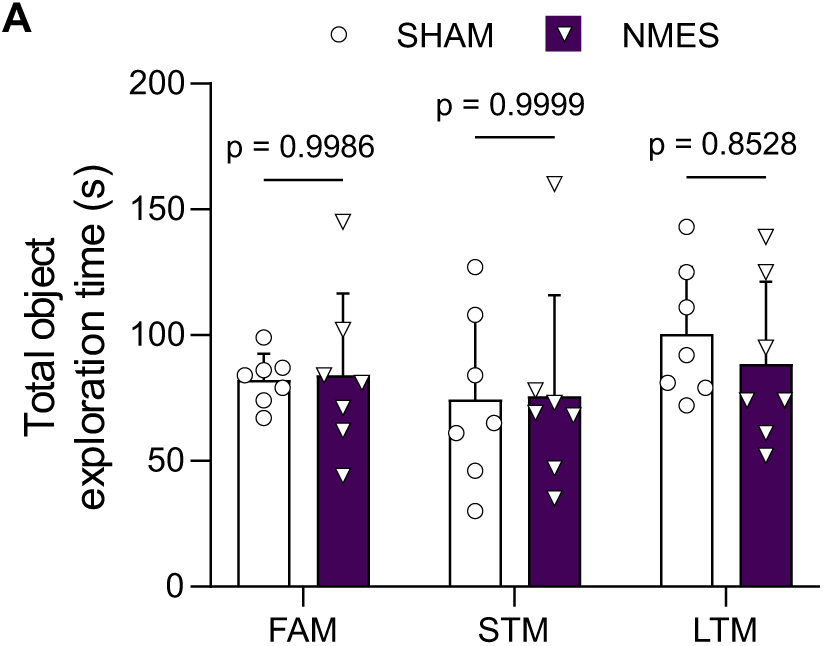
Total object exploration time during the novel object recognition task. (A) Total time spent exploring both objects during the familiarization phase (FAM), short-term memory test (STM), and long-term memory test (LTM) in SHAM-and NMES-treated animals. Data are presented as individual values with mean ± SD. Comparisons between groups were performed using the appropriate two-tailed unpaired test. No between-group differences were detected at any phase of the task.

## Notes

### Competing Interest Statement

The authors have declared no competing interest.

